# Cross-sectional association study from NHANES between some dietary supplements and anti-HAV antibodies

**DOI:** 10.64898/2025.12.29.696889

**Authors:** Hussein S. Salama, Hadeer M. Nageeb, Mostafa I. Abdelglil, Mingxuan Hou, Xi Zheng, Yangyang Dong, Ling Yang

## Abstract

**Background/Objectives:** dietary supplements intake is known for its importance in preserving immune system and immune response but we need to estimate if there were direct cross-sectional association between some dietary supplements use 24-hours and anti-HAV antibodies or not.

**Methods:** data merged from three NHANES three cycles along (2013-2018) were analyzed, representing 29.400 participants across all ages. The data collected were subjected to a logistic regression analysis to assess the relationship between the reported frequency of infections and dietary nutrients consumed including minerals and some vitamins.

**Results:** the results in overall verified no significant association between investigated dietary supplements use 24-hours and anti-HAV antibodies. However, vitamin D (p = 0.004) showed a significant positive association.

**Conclusions:** there were true direct association revealed of vitamin D on anti-HAV antibodies.

## Introduction

Globally, hepatitis A virus (HAV) considered one of the most infectious etiologies of acute hepatitis. HAV infection transmission could be done through the fecal-oral route by contaminated food, water, or close physical contact with an infectious person. The development of fulminant hepatitis is rare with HAV infection, unlike hepatitis B or C. Typical symptoms of acute infection include nausea, vomiting, abdominal pain, fatigue, malaise, poor appetite, and fever; management is with supportive care (1).

Previous studies reported the importance of HAV vaccination for children 12 months or older, adults with the risk of exposure, and chronic liver disease patients (2–4). After routine vaccination of hepatitis started in 1999, the hepatitis A incidence has declined by 90% to the ratio of 1.2 cases per 100,000 population (5). In contrast to Mexico which has a high prevalence of individuals with anti-HAV antibodies, the United States has a low prevalence. Globally, the rates of HAV have decreased due to improvements in public healthcare policies, sanitation, and education, but infection rates of other hepatitis viruses appear to be increasing After routine vaccination of hepatitis started in 1999, the hepatitis A incidence has declined by 90% to the ratio of 1.2 cases per 100,000 population (5).

Different studies reported the importance of immunomodulators as vitamins A, C, D3, E, and β-carotene as well as microelements such as zinc, selenium, iron, omega-3 fatty acids, and live active probiotic bacteria for their contribution to a stronger cellular response by increasing the number of monocytes which produce MHC II molecules (6). So, in this study we tried to studied the direct anti-HAV antibodies correlation with minerals and some vitamins supplements use 24-hour.

## Methods

### Study design and data collection

The cross-sectional analysis used publicly available data merged using the participant identifier (SEQN) from the U.S. National Health and Nutrition. Key files included: Demographics (DEMO): sex, race/ethnicity, education. Dietary intake (DS1TOT_J, DS2TOT_J): 24-hour dietary recall for minerals and some vitamins. Laboratory examination (HEPA_J): hepatitis A antibodies. Examination Survey (NHANES) spanning three continuous cycles (**cycle 1**; 2013-2014, **cycle 2;** 2015-2016, and **cycle 3;** 2017-2018). A total participant (29,400) was enrolled across the three cycles (10,175; 9,971; and 9,254, respectively). Data from three NHANES cycles (2013–20124, 2015–2016, and 2017–2018) were combined following the NHANES Analytic Guidelines. Six-year MEC weights (WTMEC6YR) were computed by dividing the 2-year examination weights (WTMEC2YR) by three, to account for the inclusion of three survey cycles. The analysis proceeded as follows. Participants were eligible if they completed the household interview and the mobile examination center visit and had available serologic testing for anti-HAV. We excluded participants who lacked anti-HAV results (n = 6397), and those with missing covariate data for age, sex, race, education, and BMI. After these exclusions, participants missing dietary supplements over 2 days were also excluded. The final analytic sample comprised n = 23003 participants.

### Nutrient variables from Individual Dietary Supplements

These variables are created by using files from the NHANES_DSD that contain information on the serving size and the quantity of each nutrient from the product label’s supplement facts panel. The participant’s reported amount taken is divided by the serving size from the label in order to determine the actual amount of nutrient consumed. The variables DS1IDS_J and DS2IDS_J indicate the actual amount of product that was consumed. The actual amount of product consumed is then multiplied by the ingredient amount for each dietary supplement or antacid.

### Anti-HAV immunoassay

Anti-HAV immunoassay estimated using a competitive immunoassay technique which involves pre-incubation of anti-HAV in the sample with HAV antigen in the assay reagent. This was followed by incubation with a conjugate reagent that contains biotinylated mouse monoclonal anti-HAV antibody and horseradish peroxidase (HRP)-labeled mouse monoclonal anti-HAV antibody. Streptavidin on the wells capture the immune complex and then the unbound materials are removed by washing (7). After that, luminescent reaction used to measure bound HRP. A reagent containing luminogenic substrates (a luminol derivative and a peracid salt) and an electron transfer agent (increases the level of light produced and prolongs its emission), is added to the wells. If the HRP conjugate with reagent antigen, it produced light due to catalyzes the oxidation of the luminol derivative. The light signals are read by the VITROS ECi/ECiQ or VITROS 3600 Immunodiagnostic System. The binding of HRP is indicative of the absence of anti-HAV antibody (8).

### Statistical Analysis

All analysis accounted for the NHANES complex survey design, incorporating MEC examination weights, along with strata and primary sampling units (PSUs), in accordance with NHANES analytics guidelines. Exp(B)for Odds Ratio (OR), significance for predicted variables, and confidence interval for OR (95% C.I. for Exp(B)) were estimated using binary logistic regression analysis using IBM SPSS statistics 22 to estimate association between dietary supplements use for 24-Hour and anti-HAV antibodies. Positive regression coefficient (B) indicates that the dependent variable (hepatitis A antibodies) increases when the predictor increase. Whereas negative regression coefficient (B) indicates that the dependent variable (anti-HAV antibodies) decreases when the predictor increase. Significance was only considered when the p-value ≤ 0.05.

Complex sample regression models including interaction terms between NHANES separate cycles and minerals uptake 24-hour were used to assess heterogeneity across survey periods. And Complex sample regression models including interaction terms between NHANES combined cycles and minerals uptake 24-hour were used to assess heterogeneity across survey periods. Plan file prepared using SDMVSTRA, SDMVPSU, and WTMEC2YR from NHANES separate and combined cycles.

### Measurement

#### 1. Dependent variable

Anti-HAV antibody status (positive/negative) was used as the dependent variable. The variable was obtained from the NHANES laboratory data file (HEPA_J) measured by a chemiluminescent immunoassay. Results were dichotomized as seropositive ( ≥ cutoff value) or seronegative (< cutoff value) according to NHANES laboratory procedures (9).

#### 2. Main exposure variables

The main exposures included daily intakes of vitamins (B1, B2, B6, B12, C, D, K, folic acid) and minerals (Ca, P, Mg, Fe, Zn, Cu, Na, K, Se) derived from the 24-hour dietary recall of dietary supplement use (DS1TOT_J and DS2TOT_J).Each variable represents the total amount (mg or µg) of nutrient intake within the 24-hour recall period for regression analysis, nutrient intake was treated as a continuous variable, and extreme outliers (values >99th percentile) were winsorized to minimize the influence of implausible reporting.

#### 3. Covariates

Covariates were selected based on prior literature and biological relevance (9). These included: Demographic factors: age (years, continuous), sex (male/female), race/ethnicity (non-Hispanic White, non-Hispanic Black, Mexican American, other Hispanic, non-Hispanic Asian, other/multiracial) — from the DEMO_J file. Socioeconomic status: education level (five categories as provided in NHANES). Health behavior factors: BMI and vaccination history.

#### 4. Variable handling and weighting

All analyses applied the NHANES complex sampling design and used the combined 6-year sample weights (WTMEC6YR) calculated according to the NHANES analytic guidelines (10). Continuous variables were summarized as means ± standard errors, and categorical variables as weighted percentages.

## Results

Results in table (1) sowed demographic characteristics as sex, race/ ethnicity, and education of U.S. participants from NHANES along three cycles (2013-2018), Cycle 1 (2013-2014); cycle 2 (2015-2016); cycle 3 (2017-2018).

**Table 1.**
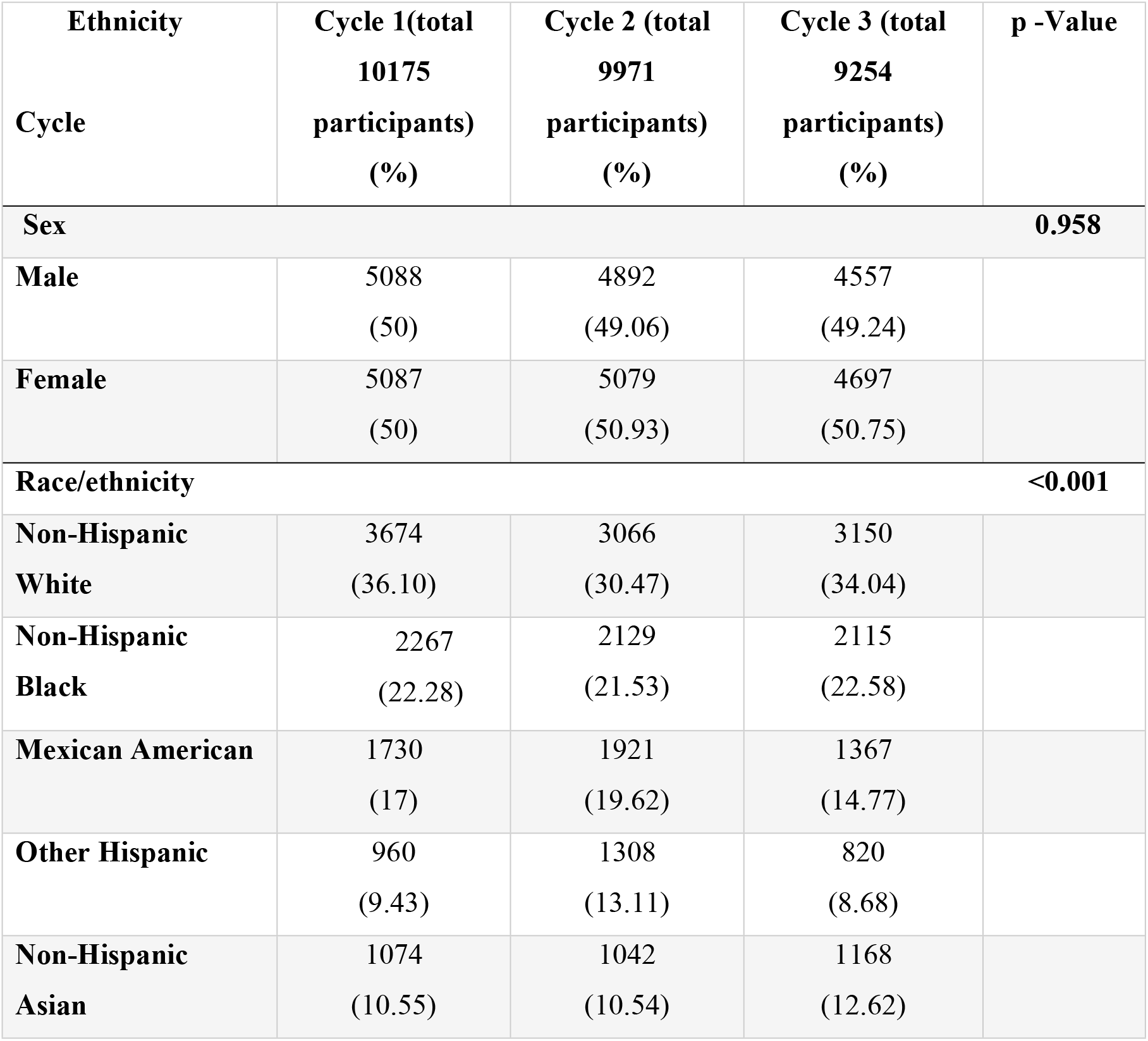
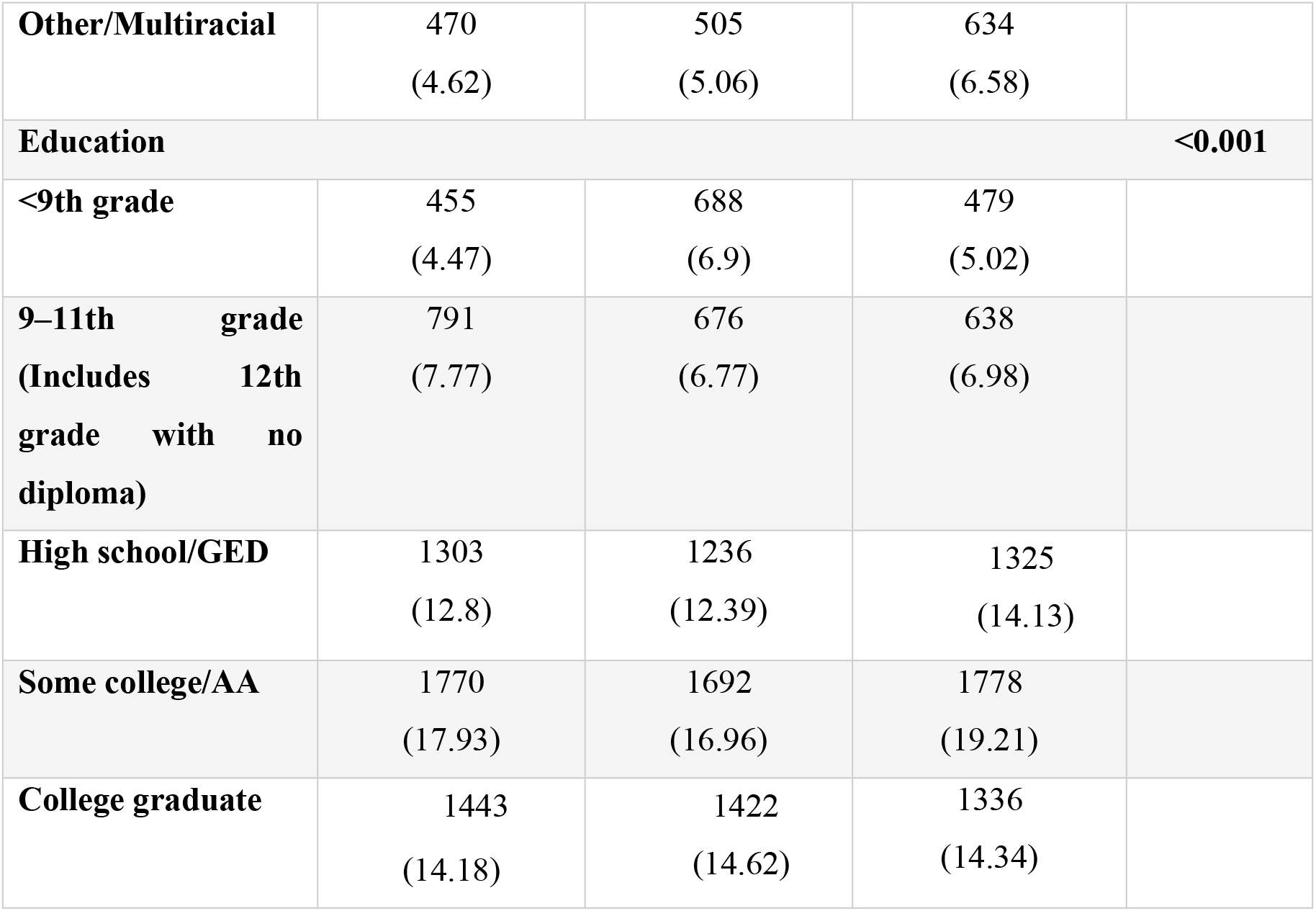
Demographic characteristics of U.S. participants (NHANES 2013-2018), Cycle 1 (2013-2014); cycle 2 (2015-2016); cycle 3 (2017-2018). A total examined participants for anti-HAV antibodies were 23003.

Table (2) sowed that, the characteristics of U.S. participants by minerals supplements use 24-hour along three cycles (2013-2018), Cycle 1 (2013-2014); cycle 2 (2015-2016); cycle 3 (2017-2018) and also the concentration of minerals supplements use 24-hour. And also, table (3) indicated the same data but on minerals supplements use 24-hour.

**Table 2.** Characteristics of U.S. participants by minerals supplements use 24-hour (NHANES 2013-2018), Cycle 1 (2013-2014); cycle 2 (2015-2016); cycle 3 (2017-2018).

| **No. of Participants** | **Cycle 1**<br>**participants**<br>**(%)** | **Cycle 2**<br>**participants**<br>**(%)** | **Cycle 3**<br>**participants**<br>**(%)** |
| --- | --- | --- | --- |
| **Minerals supplements** |  |  |  |
| **Zinc (0.0001 to 450 mg)** | 1600<br>(15.72) | 1316<br>(13.19) | 1283<br>(13.86) |
| **Calcium (0.18 to 3750 mg)** | 1579<br>(15.52) | 1302<br>(13.06) | 1267<br>(13.69) |
| **Phosphorus (0.67 to 1200 mg)** | 647<br>(6.36) | 521<br>(5.22) | 502<br>(5.42) |
| **Magnesium (0.08 to 4635 mg)** | 1205<br>(11.84) | 877<br>(8.79) | 883<br>(9.54) |
| **Iron (0.002 to 500 mg)** | 872<br>(8.57) | 682<br>(6.83) | 657<br>(7.09) |
| **Copper (0.0002 to 6.1 mg)** | 1144<br>(11.24) | 877<br>(8.97) | 866<br>(9.35) |
| **Sodium (0.1 to 1500 mg)** | 414<br>(4.06) | 378<br>(3.79) | 377<br>(4.07) |
| **Potassium (0.01 to 6750 mg)** | 722<br>(7.09) | 638<br>(6.39) | 627<br>(6.77) |
| **Selenium (0.1 to 566 mcg)** | 1001<br>(9.83) | 806<br>(8.08) | 820<br>(8.86) |

**Table 3.** Characteristics of U.S. participants by vitamins supplements use 24-hour (NHANES 2013-2018), Cycle 1 (2013-2014); cycle 2 (2015-2016); cycle 3 (2017-2018).

| **No. of Participants** | **Cycle 1**<br>participants<br>(%) | **Cycle 2**<br>participants<br>(%) | **Cycle 3**<br>participants<br>(%) |
| --- | --- | --- | --- |
| **Vitamins supplements** |  |  |  |
| **Thiamin (Vitamin B1) (0.001 to1500 mg)** | 1046<br>(10.28) | 1071<br>(10.74) | 1302<br>(14.06) |
| **Riboflavin (Vitamin B2) (0.001 to 401.7 mg)** | 1040<br>(10.22) | 1674<br>(16.78) | 1304<br>(14.09) |
| **Vitamin B6 (0.02 to 10010 mg)** | 1449<br>(14.24) | 1064<br>(10.67) | 1723<br>(18.62) |
| **Folic acid (1 to 20400 mcg)** | 1412<br>(13.87) | 1450<br>(15.44) | 1671<br>(18.06) |
| **Vitamin B12 (0.04 to 60000 mcg)** | 1553<br>(15.8) | 1562<br>(15.66) | 1760<br>(19.02) |
| **Vitamin C (0.33 to 18000 mg)** | 1607<br>(15.79) | 1635<br>(16.39) | 1876<br>(20.72) |
| **Vitamin K (0.1 to 600 mcg)** | 787<br>(7.73) | 788<br>(7.8) | 932<br>(10.07) |
| **Vitamin D (D2 + D3) (0.004 to 2570 mcg)** | 1803<br>(17.72) | 1810<br>(18.15) | 2044<br>(22.09) |

As shown in table (4) gender, age, vaccination, education, race, BMI, and all minerals showed no significance at 0.05 level. Indicating that there were no direct association between minerals intake and Anti-HAV antibodies production.

**Table 4.** Variables in the equation showing binary logistic regression analysis for minerals supplement use 24-Hour and anti-HAV antibodies.

|  |  | **B** | **S.E.** | **Wald** | **df** | **Sig.** | **Exp(B)** | **95% C.I. for EXP(B)** |  |
| --- | --- | --- | --- | --- | --- | --- | --- | --- | --- |
|  |  |  |  |  |  |  |  | **Lower** | **Upper** |
| **Step 1<sup>a</sup>** | **Gender (1)** | -.111 | .644 | .030 | 1 | .863 | .895 | .253 | 3.159 |
|  | **Age** | -.023 | .013 | 2.921 | 1 | .087 | .978 | .953 | 1.003 |
|  | **Vaccination** | .271 | .143 | 3.587 | 1 | .058 | 1.311 | .991 | 1.736 |
|  | **Race** |  |  | 3.293 | 4 | .510 |  |  |  |
|  | **Education** |  |  | 3.149 | 12 | .994 |  |  |  |
|  | **BMI** |  |  | .080 | 3 | .994 |  |  |  |
|  | **Ca** | .001 | .001 | 3.133 | 1 | .077 | 1.001 | 1.000 | 1.002 |
|  | **P** | .001 | .005 | .073 | 1 | .788 | 1.001 | .992 | 1.011 |
|  | **Mg** | -.001 | .002 | .131 | 1 | .718 | .999 | .996 | 1.003 |
|  | **Fe** | .013 | .019 | .441 | 1 | .507 | 1.013 | .976 | 1.051 |
|  | **Zn** | -.016 | .017 | .878 | 1 | .349 | .984 | .951 | 1.018 |
|  | **Cu** | .050 | .027 | 3.387 | 1 | .066 | 1.051 | .997 | 1.108 |
|  | **Na** | -.006 | .005 | 1.468 | 1 | .226 | .994 | .983 | 1.004 |
|  | **K** | .001 | .004 | .122 | 1 | .727 | 1.001 | .994 | 1.009 |
|  | **Se** | -.010 | .008 | 1.580 | 1 | .209 | .990 | .974 | 1.006 |
|  | **Constant** | 2.792 | 3.085 | .819 | 1 | .365 | 16.314 |  |  |
| **a. Variables entered on step 1: gender, age, vaccine, race, education, BMI, Ca, P, mg, Fe, Zn, Cu, Na, K, Se.** |  |  |  |  |  |  |  |  |  |

As shown in table (5), significant interactions across combined cycles were observed for phosphorus (p = 0.002), iron (p = 0.001), zinc (p < 0.001), cupper (p=.001), potassium (p < 0.001) and selenium (p < 0.001), indicating variation in these associations across cycles. No significant heterogeneity was detected for calcium, magnesium, and sodium or potassium (p > 0.05) indicating consistency of these predictors along cycles.

**Table 5.** Test of heterogeneity of minerals across cycles.

|  | Cycle 1 | Cycle 2 | Cycle 3 | Combined cycle |
| --- | --- | --- | --- | --- |
|  | Sig. | Sig. | Sig. | Sig. |
| **Ca** | .820 | .333 | .166 | .163 |
| **P** | .244 | .000* | .000* | .000* |
| **mg** | .314 | .634 | .926 | .882 |
| **Fe** | .965 | .010* | .008* | .001* |
| **Zn** | .060 | .195 | .002* | .000* |
| **Cu** | .051 | .001* | .626 | .001* |
| **Na** | .643 | .130 | .648 | .100 |
| **K** | .239 | .000* | .000* | .000* |
| **Se** | .225 | .250 | .000* | .000* |

Binary logistic regression was conducted to examine predictors of antibody presence as shown in table (6). The model included gender, age, vaccination, education, race, BMI, vitamin B1, B2, B6, B12, folic acid, vitamin C, vitamin K, and vitamin D. The overall model was statistically significant (χ^2^ = 10.937, p = 0.001). Among all predictors, only variable vitamin D showed a significant effect (B = 0.001, p = 0.004, OR = 1.001, 95% CI [1.000, 1.001]), indicating that higher values of “vitamin D” were associated with slightly increased odds of antibodies. All other predictors were not statistically significant (p > 0.05).

**Table 6.** Variables in the equation showing binary logistic regression analysis for vitamins supplement use 24-Hour and anti-HAV antibodies.

|  |  | B | S.E. | Wald | Df | Sig. | Exp(B) | 95% C.I. for EXP(B) |  |
| --- | --- | --- | --- | --- | --- | --- | --- | --- | --- |
|  |  |  |  |  |  |  |  | Lower | Upper |
| **Step 1<sup>a</sup>** | **Gender (1)** | -.089 | .097 | .839 | 1 | .360 | .915 | .756 | 1.107 |
|  | **Age** | -.001 | .003 | .082 | 1 | .775 | .999 | .994 | 1.004 |
|  | **Vaccination** | .010 | .018 | .323 | 1 | .570 | 1.010 | .975 | 1.047 |
|  | **Race** |  |  | 8.084 | 4 | .089 |  |  |  |
|  | **Education** |  |  | 23.494 | 17 | .134 |  |  |  |
|  | **BMI** |  |  | 2.327 | 3 | .507 |  |  |  |
|  | **Vitamin B1** | .003 | .003 | .989 | 1 | .320 | 1.003 | .997 | 1.008 |
|  | **Vitamin B2** | .003 | .002 | 1.776 | 1 | .183 | 1.003 | .999 | 1.007 |
|  | **Vitamin B6** | .000 | .000 | 1.377 | 1 | .241 | 1.000 | .999 | 1.000 |
|  | **Vitamin B12** | .000 | .000 | 1.268 | 1 | .260 | 1.000 | 1.000 | 1.000 |
|  | **Folic acid** | .000 | .000 | 2.119 | 1 | .146 | 1.000 | .999 | 1.000 |
|  | **Vitamin C** | .000 | .000 | .034 | 1 | .855 | 1.000 | 1.000 | 1.000 |
|  | **Vitamin K** | .001 | .001 | .604 | 1 | .437 | 1.001 | .999 | 1.003 |
|  | **Vitamin D** | .000 | .000 | 8.391 | 1 | .004* | 1.000 | 1.000 | 1.001 |
|  | **Constant** | -<br>1.089 | 1.265 | .741 | 1 | .389 | .337 |  |  |
| **a. Variables entered on step 1: gender, age, vaccination, race, education, BMI, vitamin B1, vitamin B2, vitamin B6, vitamin B12, folic acid, vitamin C, vitamin K, vitamin D.** |  |  |  |  |  |  |  |  |  |

Results in table (7) showed significant interactions across combined cycles. Results were observed for vitamin B2 (p = 0.009, 0.014) and vitamin K (p = 0.038, 0.008) indicating variation in these associations across cycles. And no significant heterogeneity was detected for vitamin B1, vitamin B6, vitamin B12, folic acid, vitamin C, and vitamin D (p > 0.05) indicating consistency of these predictors along cycles.

**Table 7.** Test of heterogeneity of vitamins across cycles.

|  | **Cycle 1** | **Cycle 2** | **Cycle 3** | **Combined Cycles** |
| --- | --- | --- | --- | --- |
|  | **Sig.** | **Sig.** | **Sig.** | **Sig.** |
| **Vitamin B1** | .571 | .492 | .374 | .492 |
| **Vitamin B2** | .009* | .014* | .327 | .014 |
| **Vitamin B6** | .078 | .275 | .637 | .275 |
| **Vitamin B12** | .484 | .812 | .511 | .812 |
| **Folic acid** | .484 | .328 | .250 | .328 |
| **Vitamin C** | .470 | .490 | .307 | .490 |
| **Vitamin K** | .156 | .038* | .008* | .038* |
| **Vitamin D** | .092 | .267 | .381 | .267 |

## Discussion

All minerals tested showed no significant effect on Anti-HAV antibodies indicating that minerals intake for 24 hours couldn’t be used as a predictor for HAV antibodies which suggests the relationship between micronutrient status and immune response to HAV may be more complex and influenced by other factors such as nutritional status, infection history, vaccination, or genetic differences in immune function. But some literature data reported that some minerals as zinc and selenium have a well-functioning immune system effect against viral infections (11).

Complex regression analysis indicating that the relationship between variables changes significantly from one cycle (or time period) to another, rather than being consistent. This indicates that the effect phosphorous, iron, cupper, potassium, and selenium size or the nature of the relationships changes meaningfully across different periods or “cycles” in the data.

Results of vitamins and anti-HAV antibodies logistic regression suggests that none of studied vitamins have a significant independent effect on the outcome except vitamin D that showed significant positive association. This results strongly agreed with(12–14) who reported the importance of calcitriol to stimulates antimicrobial activities of macrophages and monocytes through VDR-RXR signaling, which triggers the production of cathelicidins that directly influences various respiratory viruses by disrupting viral envelopes. Also, vitamin D also affects the immune system as its calcitriol active metabolite shift the production of cytokines from Th1 to Th2 (15–17).

Other studies suggest that vitamin D sufficiency may support optimal antibody-mediated immunity. However, such effects occur over weeks to months rather than hours or days and are typically observed in relation to overall vitamin D status (serum 25(OH)D), not acute short-term supplement consumption. Thus, while vitamin D may plausibly play a long-term role in shaping immune responses, the 2-day exposure measured here cannot directly explain elevations in anti-HAV IgG (16,18). So, further Longitudinal or mechanistic studies were need to confirm direct association effect of vitamin D on anti-HAV antibodies.

None significant heterogeneity across cycles in complex regression of all vitamins except vitamin B1 and vitamin K which showed significant heterogeneity indicating consistency in results over different time periods or cycles, and there is no statistically meaningful variation in how the independent variables affect the outcome variable across those cycles.

Finally, using NHANES 2013-2018 results indicated significant cross-sectional association between using vitamin D supplements for 24-hours and anti-HAV antibodies.

## Limitations

Due to the sample size or variability was insufficient to detect subtle associations, more studies with larger populations are needed to clarify these relationships. Longitudinal or mechanistic studies would also help clarify whether chronic vitamin D status influences HAV vaccine responsiveness, although no evidence supports a role for short-term exposure.

## Conclusion

In this study binary logistic regression results revealed no significance of all predictors tested with the dependent variable (anti-HAV antibodies) except vitamin D supplements which showed a significant predictors for anti-HAV antibodies. So, further investigation of longitudinal and mechanistic evidence and investigation with a larger sample needed to confirm the revealed results.

## Declarations

### Ethics approval and consent to participate

not applicable

### Consent for publication

not applicable

### Availability of data and material

all data are available with corresponding author

### Authors’ contributions

**Conceptualization**, Hussein S. Salama, Ling Yang, Hadeer M. Nageeb; **Data curation**, Hussein S. Salama, Ling Yang, Hadeer M. Nageeb;**Methodology**, Hussein S. Salama, Hadeer M. Nageeb; **Validation**, Ling Yang; Formal analysis, Hussein S. Salama; Investigation, Mostafa I. Abdelglil, Mingxuan Hou, Xi Zheng, Yangyang Dong,Ling Yang ; **Supervision**,,Ling Yang; Data curation, Mingxuan Hou, Xi Zheng, Yangyang Dong; **Writing original draft**, Hussein S. Salama, Hadeer M. Nageeb; **Funding acquisition**, not applicable; **Writing-Review & Editing**, Hussein S. Salama, Hadeer M. Nageeb, ostafa I. Abdelglil, Mingxuan Hou, Xi Zheng, Yangyang Dong,Ling Yang

## References

1. Odenwald MA, Paul S. Viral hepatitis: Past, present, and future. World J Gastroenterol [Internet]. 2022 Apr 14;28(14):1405–29. Available from: https://www.wjgnet.com/1007-9327/full/v28/i14/1405.htm

2. Brennan J, Moore K, Sizemore L, Mathieson SA, Wester C, Dunn JR, et al. Notes from the Field: Acute Hepatitis A Virus Infection Among Previously Vaccinated Persons with HIV Infection — Tennessee, 2018. MMWR Morb Mortal Wkly Rep [Internet]. 2019 Apr 12;68(14):328–9. Available from: http://www.cdc.gov/mmwr/volumes/68/wr/mm6814a3.htm?s_cid=mm6814a3_w

3. Johnson KD, Lu X, Zhang D. Adherence to hepatitis A and hepatitis B multi-dose vaccination schedules among adults in the United Kingdom: a retrospective cohort study. BMC Public Health [Internet]. 2019 Dec 15;19(1):404. Available from: https://bmcpublichealth.biomedcentral.com/articles/10.1186/s12889-019-6693-5

4. Alberts CJ, Boyd A, Bruisten SM, Heijman T, Hogewoning A, Rooijen M van, et al. Hepatitis A incidence, seroprevalence, and vaccination decision among MSM in Amsterdam, the Netherlands. Vaccine [Internet]. 2019 May;37(21):2849–56. Available from: https://linkinghub.elsevier.com/retrieve/pii/S0264410X19303822

5. Klevens RM, Miller JT, Iqbal K, Thomas A, Rizzo EM, Hanson H, et al. The Evolving Epidemiology of Hepatitis A in the United States. Arch Intern Med [Internet]. 2010 Nov 8;170(20). Available from: http://archinte.jamanetwork.com/article.aspx?doi=10.1001/archinternmed.2010.401

6. Toti E, Chen CYO, Palmery M, Villaño Valencia D, Peluso I. Non-Provitamin A and Provitamin A Carotenoids as Immunomodulators: Recommended Dietary Allowance, Therapeutic Index, or Personalized Nutrition? Cirillo G, editor. Oxid Med Cell Longev [Internet]. 2018 Jan 9;2018(1). Available from: https://onlinelibrary.wiley.com/doi/10.1155/2018/4637861

7. NCCLS. rocedures for the Handling and Processing of Blood Specimens; NCCLS; 1999. 1-56238-388–4 p.

8. Liaw YF, Yang CY, Chu CM, Huang MJ. Appearance and persistence of hepatitis A IgM antibody in acute clinical hepatitis A observed in an outbreak. Infection [Internet]. 1986 Jul;14(4):156–8. Available from: http://link.springer.com/10.1007/BF01645253

9. Wang Z, Guo H, Teng X. Diabetic retinopathy and mortality in adults with diabetes: causal mediation analysis of the role of HbA1c. Diabetes Res Clin Pract [Internet]. 2025 Nov;229:112936. Available from: https://linkinghub.elsevier.com/retrieve/pii/S0168822725009507

10. Leroux A, Di J, Smirnova E, Mcguffey EJ, Cao Q, Bayatmokhtari E, et al. Organizing and Analyzing the Activity Data in NHANES. Stat Biosci [Internet]. 2019 Jul 9;11(2):262–87. Available from: http://link.springer.com/10.1007/s12561-018-09229-9

11. Calder P, Carr A, Gombart A, Eggersdorfer M. Optimal Nutritional Status for a Well-Functioning Immune System Is an Important Factor to Protect against Viral Infections. Nutrients [Internet]. 2020 Apr 23;12(4):1181. Available from: https://www.mdpi.com/2072-6643/12/4/1181

12. Tripathi S, Tecle T, Verma A, Crouch E, White M, Hartshorn KL. The human cathelicidin LL-37 inhibits influenza A viruses through a mechanism distinct from that of surfactant protein D or defensins. J Gen Virol [Internet]. 2013 Jan 1;94(1):40–9. Available from: https://www.microbiologyresearch.org/content/journal/jgv/10.1099/vir.0.045013-0

13. Barlow PG, Svoboda P, Mackellar A, Nash AA, York IA, Pohl J, et al. Antiviral Activity and Increased Host Defense against Influenza Infection Elicited by the Human Cathelicidin LL-37. Kovats S, editor. PLoS One [Internet]. 2011 Oct 21;6(10):e25333. Available from: https://dx.plos.org/10.1371/journal.pone.0025333

14. Sharma OP. Hypercalcemia in granulomatous disorders: a clinical review. Curr Opin Pulm Med [Internet]. 2000 Sep;6(5):442–7. Available from: http://journals.lww.com/00063198-200009000-00010

15. Polak E, Stę pień AE, Gol O, Tabarkiewicz J. Potential Immunomodulatory Effects from Consumption of Nutrients in Whole Foods and Supplements on the Frequency and Course of Infection: Preliminary Results. Nutrients [Internet]. 2021 Apr 1;13(4):1157. Available from: https://www.mdpi.com/2072-6643/13/4/1157

16. Sanlier N, Guney-Coskun M. Vitamin D, the immune system, and its relationship with diseases. Egypt Pediatr Assoc Gaz [Internet]. 2022 Oct 17;70(1):39. Available from: https://epag.springeropen.com/articles/10.1186/s43054-022-00135-w

17. Daryabor G, Gholijani N, Kahmini FR. A review of the critical role of vitamin D axis on the immune system. Exp Mol Pathol [Internet]. 2023 Aug;132–133:104866. Available from: https://linkinghub.elsevier.com/retrieve/pii/S0014480023000175

18. Girsang RT, Rusmil K, Fadlyana E, Kartasasmita CB, Dwi Putra MG, Setiabudiawan B. Correlation Between Vitamin D Status and HBsAg Antibody Levels in Indonesian Adolescents Immunised Against Hepatitis B. Int J Gen Med [Internet]. 2023 Nov;Volume 16:5183–92. Available from: https://www.dovepress.com/correlation-between-vitamin-d-status-and-hbsag-antibody-levels-in-indo-peer-reviewed-fulltext-article-IJGM

